# A Role for Nup153 in Nuclear Assembly Reveals Differential Requirements for Targeting of Nuclear Envelope Constituents

**DOI:** 10.1101/2022.05.27.493435

**Authors:** Dollie LaJoie, Ayse M. Turkmen, Douglas R. Mackay, Christopher C. Jensen, Vasilisa Aksenova, Maho Niwa, Mary Dasso, Katharine S. Ullman

## Abstract

Assembly of the nucleus following mitosis requires rapid and coordinate recruitment of diverse constituents to the inner nuclear membrane. We have identified an unexpected role for the nucleoporin Nup153 in promoting the continued addition of a subset of nuclear envelope proteins during initial expansion of nascent nuclei. Specifically, disrupting the function of Nup153 interferes with ongoing addition of B-type lamins, lamin B receptor (LBR), and SUN1 early in telophase, after the nuclear envelope (NE) has initially enclosed chromatin. In contrast, effects on lamin A and SUN2 were minimal, pointing to differential requirements for the ongoing targeting of nuclear envelope proteins. Further, distinct mis-targeting phenotypes arose among the proteins that require Nup153 for NE targeting. Thus, disrupting the function of Nup153 in nuclear formation reveals several previously undescribed features important for establishing nuclear architecture: 1) a role for a nuclear basket constituent in ongoing recruitment of nuclear envelope components, 2) two functionally separable phases of nuclear envelope formation in mammalian cells, and 3) distinct requirements of individual nuclear envelope residents for continued targeting during the expansion phase of NE reformation.

## Introduction

Metazoan cells undergo an open mitosis in which the nuclear membranes, nuclear lamina, and nuclear pore complexes (NPCs) disperse prior to partitioning of the genome. Soon after chromosomes begin to segregate, the process of nuclear assembly begins to take place around each mass of chromosomes, termed a chromatin disk. The mitotic endoplasmic reticulum (ER), which accommodates membrane proteins that are residents of the nuclear envelope (NE), begins to contact and enclose the chromosomes at early anaphase (LaJoie and Ullman, 2017; Schellhaus et al., 2016). At this time, the inner and outer membranes of the NE are established and a diverse repertoire of NE constituents target to the chromatin surface, with some taking on roles critical to coordinating nuclear assembly. One such role is recruitment of machinery to synchronously remodel microtubules and seal membranes, which takes place at subdomains within the nuclear envelope termed ‘core’ regions. Core regions form transiently in the center of anaphase disks, on both inner and outer faces, with the remainder of the NE termed the ‘non-core’ region (Liu and Pellman, 2020). Members of the endosomal sorting complexes required for transport (ESCRT) pathway are recruited to this site and play a central role in sealing the NE in a coordinate fashion (Stoten and Carlton, 2018). Concomitantly, rapid nuclear pore formation takes place at the non-core region (Otsuka and Ellenberg, 2018). Thus, once the NE is sealed, the nucleus is competent to establish specialization of the nuclear environment via nucleocytoplasmic transport. To fully form normal nuclear architecture, however, the NE must also continue to expand. Indeed, coordinated lipid synthesis in conjunction with ESCRT-mediated sealing was found to be important in nuclear assembly (Penfield et al., 2020). At the transition from closure to expansion, NE proteins redistribute throughout the INM, removing the distinction between core and non-core regions (Maeshima et al., 2006). NPCs continue to be added, but their biogenesis at this stage, which is termed an ‘interphasic’ or ‘inside-out’ route, has distinct features due to the necessity to form this macromolecular structure in what is now a contiguous double lipid bilayer (Otsuka and Ellenberg, 2018).

Beyond this second, distinct route of NPC formation, for which an understanding has begun to emerge (Doucet and Hetzer, 2010; Dultz and Ellenberg, 2010; Otsuka et al., 2016; Otsuka et al., 2021), little is known about the second phase of nuclear assembly, NE expansion. Early work in the *Xenopus* egg extract system identified roles for the AAA-ATPase p97, mediated by specific co-factors that were key in defining distinct phases of NE sealing versus continued nuclear growth (Hetzer et al., 2001). This has since been appreciated to reflect a role for p97 in removing Aurora B associated with chromatin (Dobrynin et al., 2011), as well as in establishing a reticular ER network that efficiently encloses chromatin (Hetzer et al., 2001). ER shaping proteins actively regulate ER morphology during nuclear assembly to promote proper reformation of the nuclear envelope at a time when INM proteins are being targeted to and enriching at the chromatin surface (Anderson and Hetzer, 2008; Lu et al., 2009, 2011). It is of note that the architecture of the endoplasmic reticulum, and particularly the role of Atlastin, is established to be important for INM protein diffusion through the ER and subsequent targeting to the NE at interphase (Pawar et al., 2017). There has also been considerable focus on how proteins target to the INM during ongoing recruitment of newly-synthesized INM proteins at interphase (Boni et al., 2015; King et al., 2006; Lokareddy et al., 2015; Meinema et al., 2011; Ohba et al., 2004; Rempel et al., 2020; Ungricht et al., 2016). These studies point toward pathways for INM protein passage through the NPC, after which interactions at the nuclear periphery may aid in retention. Nuclear size control has additionally been scrutinized, reinforcing the importance of perinuclear ER and revealing several transport related factors that dictate the extent and maintenance of nuclear dimensions (Jevtic et al., 2019; Mukherjee et al., 2020; Vukovic et al., 2016). Although post-mitotic NE expansion may rely on some of the same mechanisms studied in these other contexts, the results we present here reveal requirements in mammalian cells unique to NE growth during initial nuclear formation. Moreover, our results also expose distinctions among INM proteins in terms of their requirements for NE targeting at this stage.

Failure to form a normal nuclear envelope environment results in impaired compartmentalization and DNA damage (Vietri et al., 2015; von Appen et al., 2020). This is exemplified by micronuclei, which are hallmarks of tumor cells and form from chromosomes aberrantly separated from the primary nucleus due to missegregation events. Nuclear envelope formed at these sites often fails to recruit the full repertoire of INM proteins and is similar to the core domain at the newly-forming primary nucleus: enriched in LEM-domain proteins and deficient in B-type lamins and NPCs (Liu et al., 2018). Such micronuclei suffer from unrestrained membrane remodeling events by the ESCRT-III pathway (Vietri et al., 2020; Willan et al., 2019) and are prone to rupture (Hatch et al., 2013). Ultimately micronuclei experience increased DNA damage, conferring genome instability to an already vulnerable chromosome or chromosomal fragment (Hatch et al., 2013; Liu et al., 2018; Zhang et al., 2015).

This report focuses on Nup153, a constituent of the nuclear pore that is part of a basket-like structure emanating from the nuclear face of the NPC (Ball and Ullman, 2005). We have found that Nup153 plays a novel role in promoting nuclear assembly that is required for targeting B-type lamins and a subset of inner nuclear membrane proteins during telophase. The timing of this role uncouples early targeting events from the continued addition of certain nuclear envelope constituents. These data reveal unexpected and distinct NE protein targeting requirements and highlight the existence of a specific phase of nuclear formation.

## Results

### Nup153 plays a role in lamin B localization after mitosis

We previously reported that Nup153 depletion results in ectopic localization of lamin proteins in newly divided cells. Cells with this phenotype remain attached by an intercellular bridge containing an elongated microtubule midbody structure but otherwise appear to have interphase nuclear morphology, indicating they are in late telophase and have recently undergone nuclear assembly (Mackay et al., 2010). To further assess the role of Nup153 in nuclear lamina formation, we employed a panel of antibodies that discriminate between lamins A/C, B1, or B2 (Supplemental Figure 1). Using these isoform-specific antibodies, we found lamin A/C to be weakly mistargeted when Nup153 is depleted, whereas lamins B1 and B2 had a robust ectopic localization phenotype in late telophase cells (Figure 1A). In fixed cell analysis, endogenous lamin B2 mistargeting in Nup153-depleted cells was detectable in some early telophase cells (identified by a compact midbody structure connecting cells that have not fully flattened), with a robust phenotype present in 80±13% of cells at late telophase (Figure 1B). These observations suggested a role for Nup153 in the continued addition of B-type lamins after early targeting events in nuclear formation have completed. Alternatively, this phenotype could arise from an autophagic process of lamin B removal similar to what has been observed during senescence (Dou et al., 2015). To address these possibilities, we monitored the development of the phenotype in real-time. Live imaging of cells confirmed that early steps of nuclear assembly proceed normally in cells depleted of Nup153. Specifically, a canonical ‘core’ domain at anaphase marked by LEM2-mCherry formed while GFP-lamin B2 was recruited to non-core regions and subsequently encircled the chromatin disk similarly in both siControl and siNup153-treated cells (Supplemental Figure 2). In cells co-expressing H2B-mCherry and GFP-lamin B2, normal lamin B2 targeting to the chromatin surface during anaphase was evidenced by GFP-lamin B2 coating the nuclear surface at the time of complete cleavage furrow ingression (defined here as t=0’). However, at the transition from anaphase to telophase, lamin B2 initiated accumulation at focal sites in the cytoplasm in cells depleted of Nup153 (Supplemental Figure 2, Figure 1C). The appearance of ectopic lamin B2 at sites distant from the nucleus supports the conclusion that lamin B is not being removed from the nuclear surface, but rather is not targeting normally to the nuclear surface once the chromatin is enclosed by membrane.

**Figure 1.**
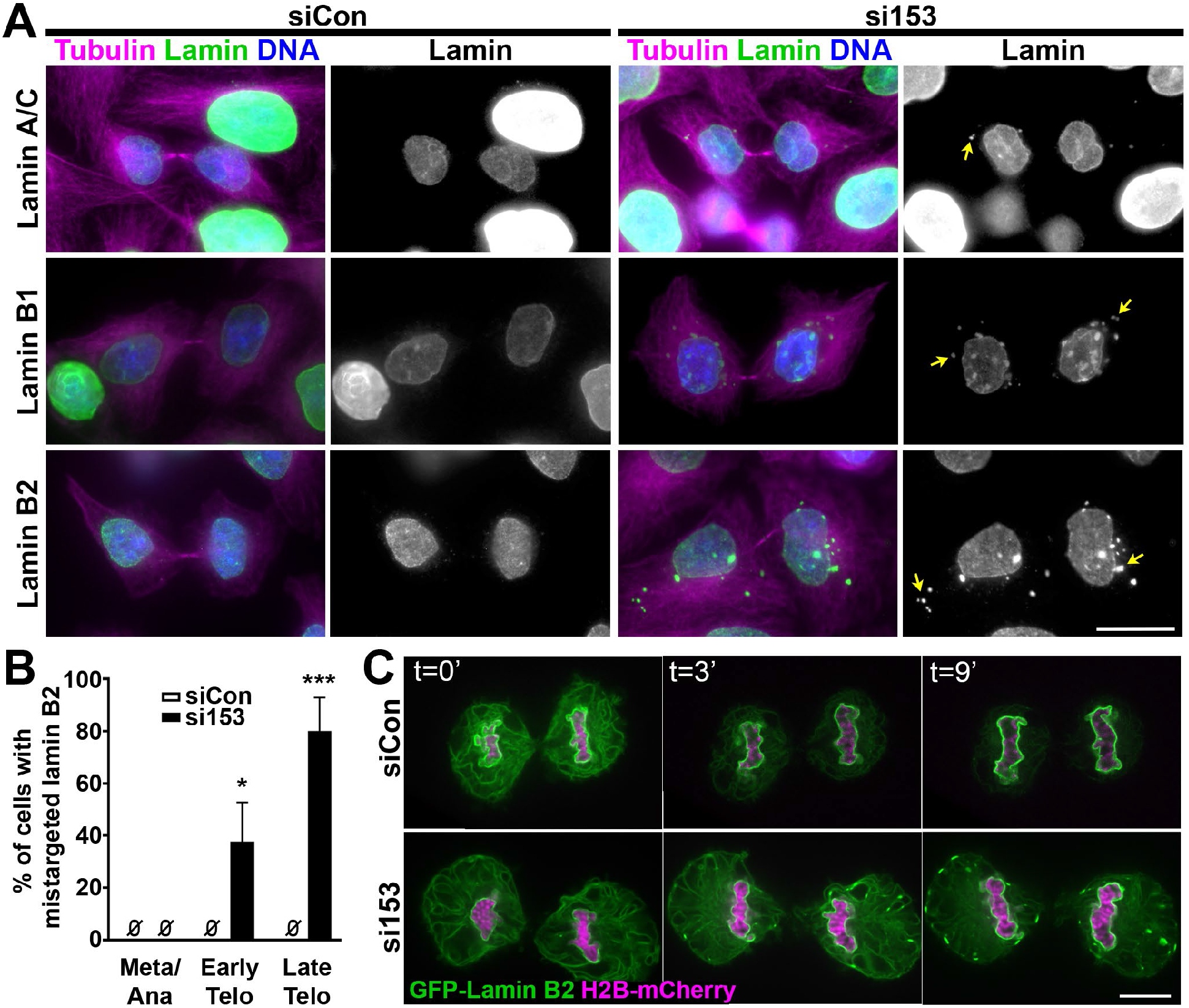
Lamins B1/B2 accumulate at ectopic sites in telophase when Nup153 is depleted. (A) Widefield microscopy showing representative images of lamins A/C, B1 and B2 in late telophase stage cells, after treatment with siControl (siCon) or siNup153 (si153) oligos for 48 hours. Yellow arrows indicate sites of lamin ectopic targeting. Scale bar 20 µm. (B) Quantification of the percent of cells with endogenous lamin B2 at metaphase/anaphase (Meta/Ana), early telophase (Early Telo), and late telophase (Late Telo) after 48-hour treatment with siCon or si153 oligos. The phenotype observed in early telophase cells was notably weaker. Data from three experiments plotted as mean ± SD; siCon: meta/ana 0±0%, early telo 0±0%, late telo 0±0%; si153: meta/ana 0±0%, early telo 38±15%, late telo 80±13%; *P < 0.05, ***P < 0.001. Ø indicates raw values were equal to 0. (C) Montage of cells expressing GFP-lamin B2 and H2B-mCherry progressing through telophase captured by live spinning disk microscopy after treatment with siCon or si153 for 48 hours, with t=0’ being the time of complete cleavage furrow ingression observed by time-lapse video. Images were adjusted independently to optimize visibility of the markers. Scale bar 10 µm.

Treatment with an independent siRNA directed against Nup153 similarly resulted in ectopic targeting of lamin B2 (si153-1, Supplemental Figure 3A), with reduction of Nup153 protein levels following siRNA treatment corroborated by immunofluorescence and immunoblot for both oligos (Supplemental Figure 3A-B). Moreover, non-transformed retinal pigment epithelial (RPE-1) cells also exhibit ectopic lamin B2 mistargeting in late telophase cells when Nup153 levels are depleted (Supplemental Figure 3C). Further substantiating our findings, induced expression of the C-terminal domain of Nup153 (Nup153-C), which we previously found to behave in a dominant negative fashion (Mackay et al., 2010), also resulted in lamin B2 mistargeting in Hela cells (Supplementary Figure 3D).

As Nup153 is known to be critical for localization of the nuclear pore basket proteins Tpr and Nup50 in late telophase cells (Aksenova et al., 2020; Hase and Cordes, 2003; Mackay et al., 2010), we sought to distinguish whether the lamin phenotype observed was due to displacement of these basket constituents or to disruption of a distinct role of Nup153. We found that neither depletion of Tpr nor depletion of Nup50 (Figure 2A) altered lamin B2 targeting (Figure 2B-C), indicating that the observed role for Nup153 function in lamin targeting is unique and not due to the absence of other constituents of the nuclear pore basket.

**Figure 2.**
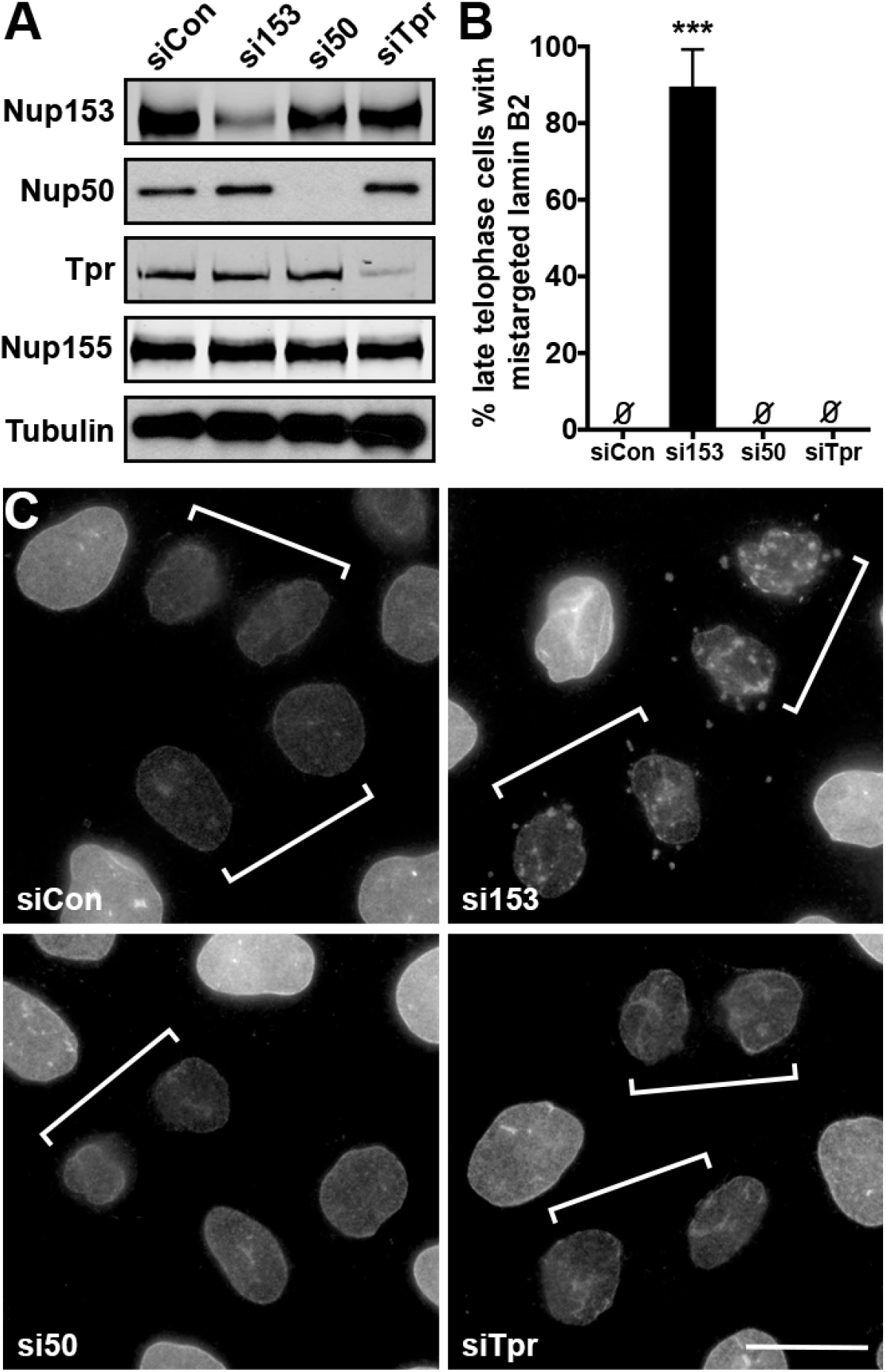
Lamin B2 mistargeting is not the result of disrupting the pore basket. (A) Immunoblot of Nup153, Nup50, Tpr, Nup155 and tubulin protein levels after depletion with indicated siRNAs for 48 hours. Nup155 and tubulin were probed as nucleoporin and loading controls, respectively. (B) Quantification of the percent of late telophase cells with mistargeted lamin B2 after treatment with the indicated siRNAs. Data from three experiments plotted as mean ± SD; siCon: 0±0%; si153: 90±10%; si50: 0±0%; siTpr: 0±0%; ***P < 0.001. Ø indicates raw values were equal to 0. (C) Widefield microscopy of lamin B2 for each depletion condition; brackets indicate late telophase cells determined by tubulin co-stain (not shown). Scale bar 20µm.

### Disrupting Nup153 functions reveals different targeting requirements for nuclear envelope proteins

We previously reported that depleting Nup153 results in an Aurora B-dependent delay in abscission. Although this prolongs the duration of cytokinesis, only ∼15% of cells are at the post-mitotic stage where this nuclear assembly phenotype manifests in Nup153-depleted cell cultures (Mackay et al., 2010). Thus, to further elucidate the role of Nup153 in nuclear formation, we sought a system more amenable to studying events at this cell cycle juncture quantitatively. To this end, we treated cells with thymidine to block cell division in G1/S followed by wash-out to release. Using this method, we found that, as expected, the number of late telophase cells, tracked by the appearance of phospho-Aurora B at an intercellular bridge of tubulin (yellow arrows), was similar in siControl- and siNup153-treated cells 10 hours post-release (white arrows indicate early telophase cells, Supplemental Figure 4). By 16 hours post-release, however, siControl-treated cells had progressed through cytokinesis while 46±4% of Nup153-depleted cells had accumulated in late telophase, where cell cycle progression is paused due to the abscission checkpoint. Under these conditions, 81±10% of late telophase cells depleted of Nup153 exhibited ectopic lamin B2 targeting, a proportion consistent with asynchronous experiments (80±13%; Supplemental Figure 4D and Figure 1D). Therefore, this strategy enriches for late telophase cells with an unaltered incidence of lamin B2 mis-targeting following Nup153 reduction, providing an ideal system in which to quantify additional aspects of this phenotype.

The selectivity of the lamin-targeting phenotype to B-type lamins, which – unlike lamin A – stably associate with membrane via a farnesyl moiety, prompted us to probe whether other proteins associated with NE membrane also mistarget when Nup153 levels are reduced. Two hallmark transmembrane proteins of the NE are SUN1 and SUN2, which each make contact with the nuclear lamina juxtaposed on the INM while interacting with outer nuclear membrane KASH-domain containing proteins in the NE lumen (Crisp et al., 2006). Similar to lamin B2, SUN1 robustly mistargeted in late telophase cells when Nup153 function was disrupted either by siRNA depletion or following expression of the dominant negative C-terminal fragment of Nup153 (Figure 3A, Supplemental Figure 5A). Given this result, it was surprising to find that SUN2 did not display notable mistargeting in Nup153-depleted cells at late telophase (Figure 3B). SUN1 mistargeting to ectopic sites occurred in a similar proportion of late telophase cells as observed for lamin B2, with SUN1 enriched 2-fold at ectopic sites relative to the NE (Figure 3C-D). Assessing SUN2 at the sites of ectopic SUN1 enrichment revealed SUN2 was not excluded from these sites, however it was present below levels found at the nucleus (Figure 3D). Plot line profile analysis of spinning disk confocal images further corroborated the conclusion that SUN1 but not SUN2 was significantly mistargeted when Nup153 levels are reduced (Figure 3E-F).

**Figure 3.**
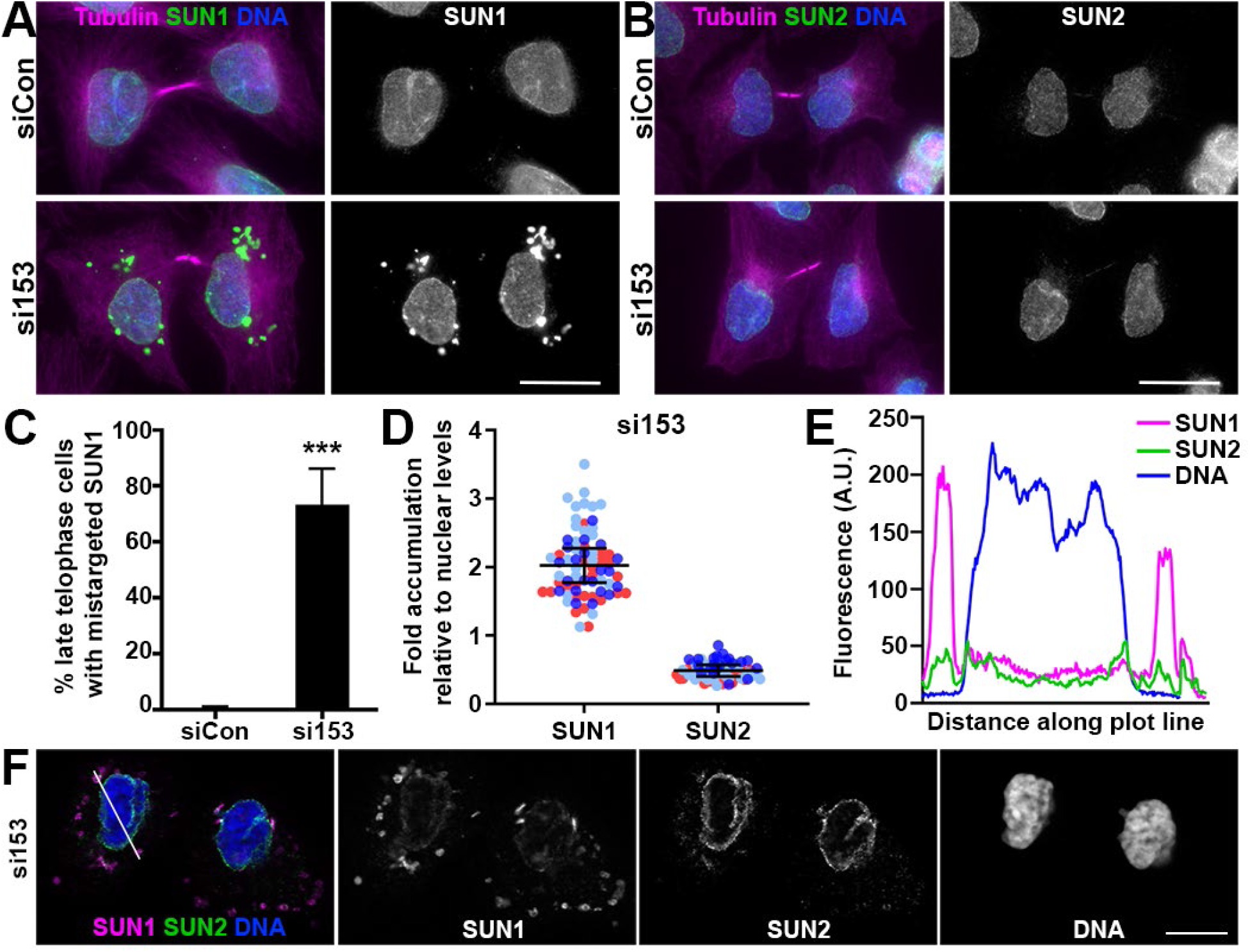
SUN1, but not SUN2, enriches at ectopic sites in late telophase cells when Nup153 is depleted. (A) Widefield microscopy of SUN1 and tubulin and SUN2 and tubulin (B) in late telophase cells following treatment with siCon or si153 under thymidine-synchronization conditions. Scale bar 20 µM. (C) Quantification of the percent late telophase cells with ectopic SUN1 targeting. Data from three experiments plotted as mean ± SD; siCon: 1±1%; si153: 73±13%; ***P < 0.001. (D) Relative levels of SUN1 and SUN2 at ectopic foci compared to nuclear levels under si153 conditions. Data collected from three independent biological replicates are depicted in different colors, N=25, 26, 20; the mean of each experiment was used to calculate the mean ± SD, indicated in black; SUN1: 2.0± 0.3%; SUN2: 0.5± 0.1%. (E) Graphed arbitrary fluorescence of SUN1 and SUN2 along plot line profile indicated on the spinning disk confocal micrographs shown in (F). Scale bar in F, 10 µm.

### Nuclear envelope proteins co-localize at ectopic NE-like membrane domains distinct from micronuclei

To gain further insight into the phenotype of aberrant targeting of particular NE residents when Nup153 function is compromised, we further characterized the focal cytoplasmic accumulation of mis-targeted proteins. We first asked the question of whether lamin B1 and B2 co-localize and found that antibodies against these B-type lamins co-stained ectopic foci when Nup153 is depleted (Figure 4A). Consistent with our finding that the incidence of ectopic lamin B2 and SUN1 accumulation was similar in late telophase cells following Nup153 depletion (Figure 3C, Supplemental Figure 4D), we found that these proteins also co-enriched at ectopic sites when assessed by spinning disk confocal microscopy (Figure 4B). Plot line profile analysis confirmed that both SUN1 and lamin B2 concentrated at these ectopic sites relative to their levels at the nucleus. In assessing additional INM residents, we found that Lap2β co-localized robustly at these cytoplasmic sites, but in contrast to SUN1, retained normal levels of targeting to the nuclear rim (Figure 4D,E).

**Figure 4.**
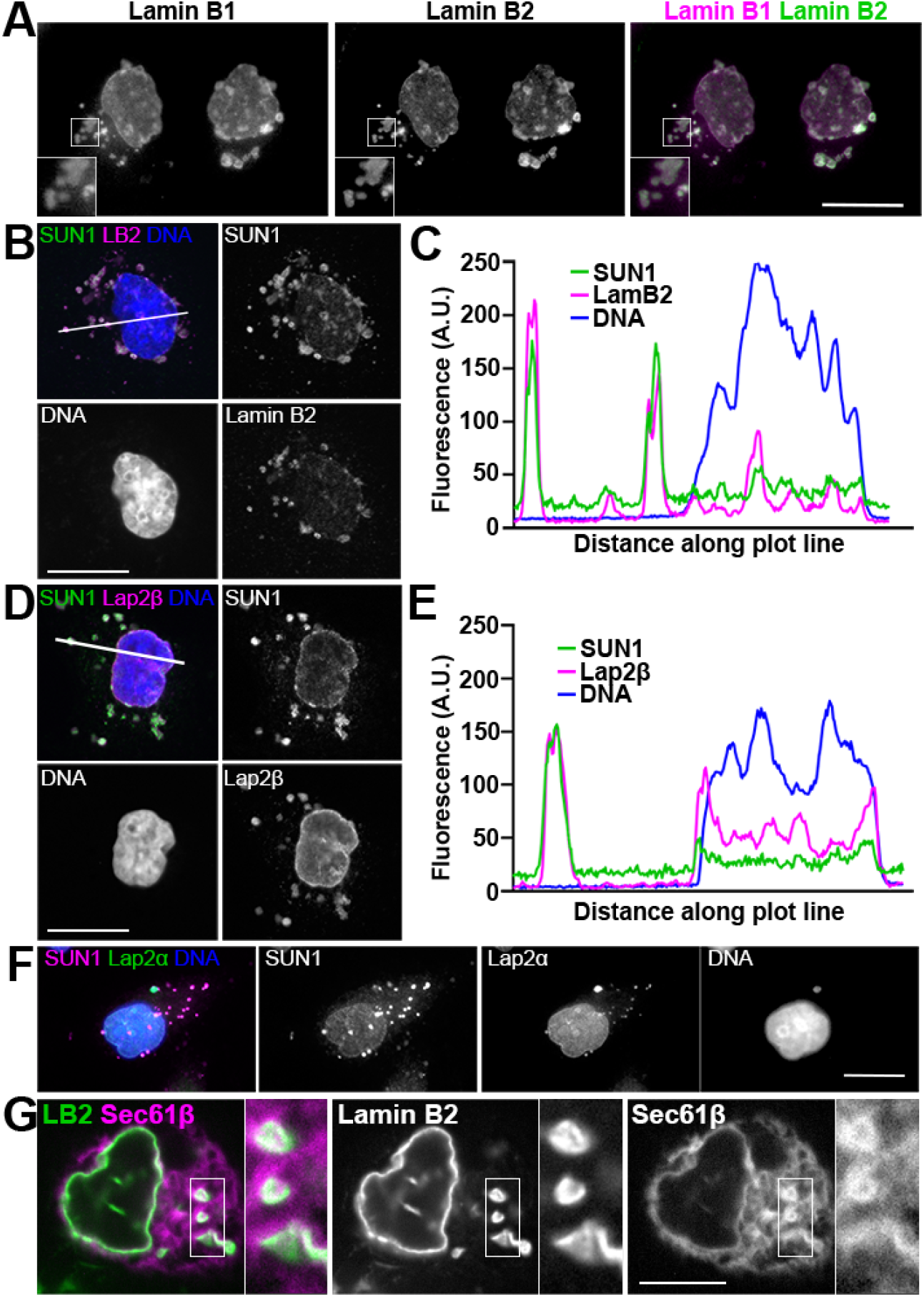
Mistargeted nuclear envelope proteins co-enrich at ectopic sites that are membranous but distinct from micronuclei. (A) Representative wide-field images of lamin B1 and B2 in Nup153-depleted cells after thymidine-synchronization. Enlargements are 2x; Scale bar 20 µm. (B) Spinning disk confocal micrographs showing lamin B2 and SUN1 after Nup153 depletion. Scale bar 20µm. (C) Arbitrary fluorescence units corresponding to plot line shown in B are graphed. (D) Spinning disk confocal micrographs showing Lap2β and SUN1 after Nup153 depletion. Scale bar 20µm. (E) Arbitrary fluorescence units corresponding to plot line shown in D are graphed. (F) Co-stain of SUN1 and Lap2α in Nup153-depleted cells by widefield microscopy, distinguish micronucleus from sites of SUN1 accumulation. Scale bar 20µm. (G) Live spinning disk confocal microscopy of BFP-Sec61β and GFP-lamin B2 in si153-treated cell, showing continuity of ER membrane with ectopic site of lamin B2 localization. Enlargements are 2.5x. Scale bar 10µm.

Despite the accumulation of membrane-associated NE proteins, DNA was not detected at these ectopic sites, as illustrated by plot line profile analysis of spinning disk confocal micrographs (Figure 3F; Figure 4C,E). This indicates that the aberrant accumulation of NE constituents following Nup153-depletion is not due to missegregation and subsequent micronuclear formation. Further reinforcing this conclusion, the nuclear protein Lap2α was found to enrich at micronuclei as expected (Liu et al., 2018), while present only at low levels at ectopic NE-like domains resulting from Nup153 depletion (Figure 4F). Indeed, while micronuclei often lack constituents such as B-type lamins and are enriched in lamin A and LEM-domain proteins (de Castro et al., 2018; Liu et al., 2018), the NE-like foci that form upon Nup153-depletion are instead enriched with proteins that are frequently low or absent at micronuclei (Figure 4D). Probing SUN1 and SUN2 at micronuclei, we found that SUN2, but not SUN1, enriched at micronuclei marked by high levels of LEM2 (Supplemental Figure 6). This pattern again contrasts with that seen following Nup153 depletion, which altered SUN1, not SUN2, targeting to the NE.

Although the sites of NE protein accumulation that result from interference with Nup153 function are not micronuclei, the presence of integral membrane proteins suggested that these focal domains are membranous. As NE proteins intermix with the endoplasmic reticulum at mitosis (Ellenberg et al., 1997; Yang et al., 1997), we expressed BFP-Sec61β and tracked its appearance in live cells. This canonical ER marker was found at sites of ectopic lamin B2 accumulation seen after Nup153 depletion, demonstrating continuity with the endoplasmic reticulum membrane, similar to its continuity with the NE proper (Figure 4G). This is also consistent with the appearance of lamin B2 in live-imaging (Figure 1C, Supplemental Figure 2), where this marker was seen in a reticular pattern in both control and Nup153-depleted cells, consistent with the morphology of the mitotic ER, and then accumulated at ER-associated sites in Nup153-depleted cells.

### A core/non-core framework reveals further complexity in targeting requirements of NE proteins

Although not originally anticipated, the pattern of Nup153-dependency in ongoing targeting of NE proteins appeared to reveal a requirement for Nup153 particular to non-core NE proteins (SUN1 and lamin B vs. SUN2, lamin A, LEM2). Targeting of LAP2β, a protein that has been detected at the non-core region (Dechat et al., 2004), was also dependent on Nup153, although its distribution was not altered to the same extent. Since NPC assembly occurs concomitantly with the early events of nuclear assembly in the non-core region, we probed the localization of POM121, an integral membrane protein of the nuclear pore, in late telophase cells. Following interference with Nup153 activity either via siRNA depletion or expression of a dominant negative fragment, late telophase cells with ectopic targeting of lamin B2 show no ectopic targeting of POM121 (Supplemental Figure 5B-D). As POM121 is first recruited to the non-core region, at face value this unperturbed localization does not fit the framework of a connection between Nup153 and non-core protein targeting. Yet, it is consistent with assembly of NPCs via a Nup153-independent ‘post-mitotic’ route at this site (Vollmer et al., 2015). Recruitment of NPC components may operate independently of requirements that distinguish core/non-core proteins even though pore formation is spatially restricted to the non-core region initially.

To further probe the core/non-core context of Nup153-dependency, we tracked another non-core NE protein, Lamin B Receptor (LBR). In this case, there was a clear dependency on Nup153 for LBR NE targeting: LBR accumulated at the NE less efficiently when Nup153 was impaired and instead a significant population was detected throughout the endoplasmic reticulum in late telophase cells (Figure 5A-B). Curiously, LBR did not greatly enrich at ectopic sites that include SUN1, although it is not excluded when assessed by plot line profile analysis following confocal microscopy (Figure 5C-D). This phenotype is similar to that observed at interphase by Mimura et al. who also demonstrated that Nup153 is critical for de-phosphorylation of LBR in interphase, which in turn promotes its proper localization at the nuclear rim (Mimura et al., 2016). A similar mechanism may be employed during initial stages of nuclear formation. Here, the significance is that Nup153 appears important to the incorporation of many non-core NE proteins during a distinct phase of NE formation and, further, targeting heterogeneity is exposed by the different phenotypes elicited for specific NE proteins when Nup153 function is impaired. The reliance on Nup153 for targeting of specific NE residents instead reveals an unanticipated role for this nucleoporin in shaping the NE repertoire, and the distinct phenotypes observed (i.e., focal concentration at NE-like domains or ER-wide dispersal) reveal additional complexity in protein trafficking requirements of individual INM residents.

**Figure 5.**
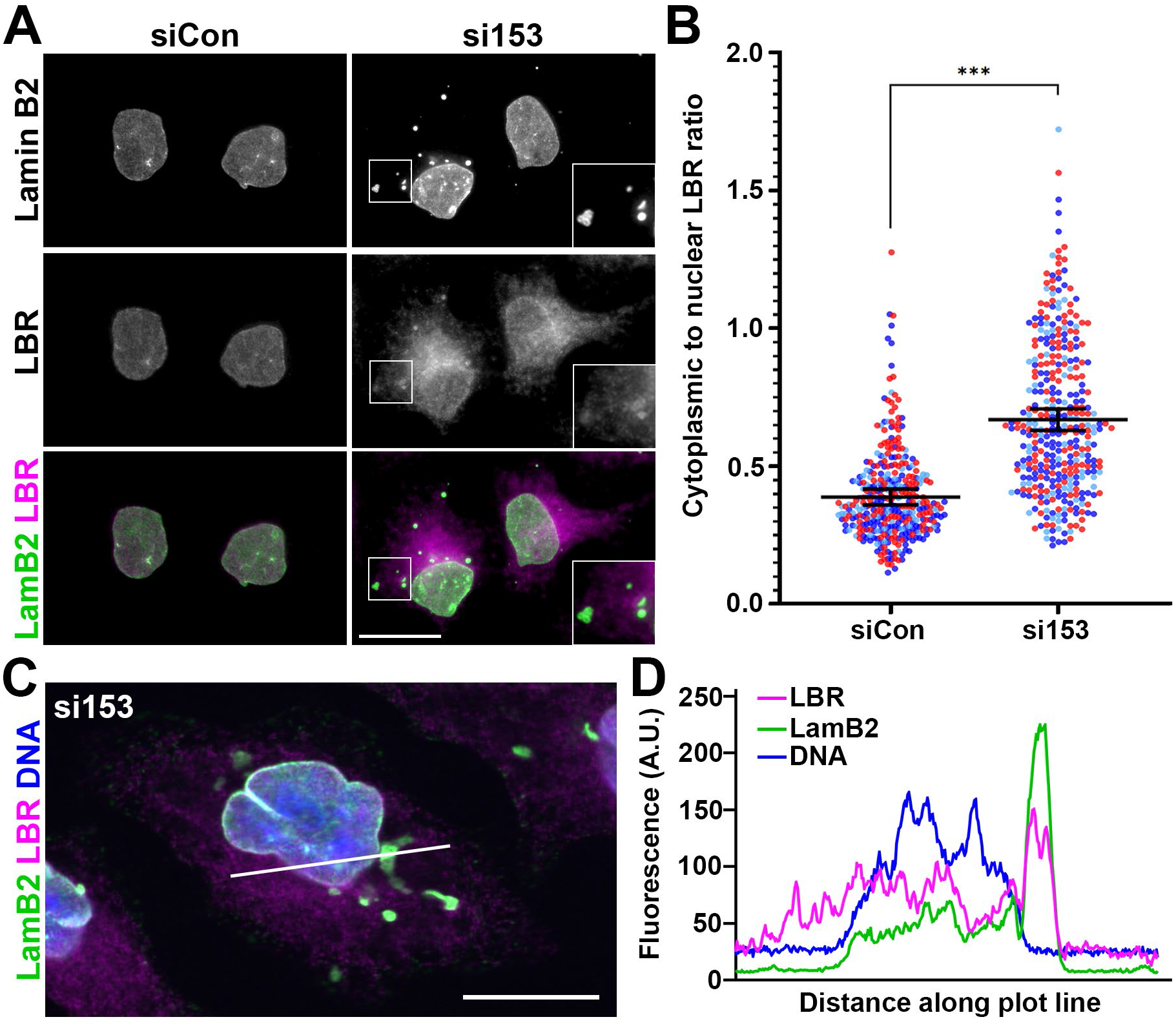
LBR accumulates ectopically in late-telophase stage cells depleted of Nup153. (A) Widefield images of LBR and lamin B2 detection in late-telophase cells, determined by tubulin staining (not shown), following treatment with siCon and si153 oligos. Scale bar is 20 μm. (B) Quantification of cytoplasmic to nuclear ratio for LBR. Data collected from three biological replicates are depicted in different colors, N =71, 68, 36 (siCon) and 72, 69, 36 (si153). The mean of each experiment was used to calculate the mean ± SD, indicated in black; siCon: 0.39 ± 0.03; si153: 0.67 ± 0.04. (C) Confocal image of LBR and lamin B2 co-detection in siNup153-treated cells. Scale bar, 20 µM. (D) Arbitrary fluorescence units corresponding to the plot line shown in C are graphed.

### Nup153 is required to establish a normal INM protein repertoire specifically during nuclear assembly

To determine whether dependence on Nup153 for protein targeting to the NE is restricted to newly forming nuclei, we employed an inducible degradation system to control Nup153 levels (Aksenova et al., 2020). This technique allows for rapid depletion, enabling the assessment of whether NE phenotypes require passage through mitosis. Using DLD-1 cells, a colorectal adenocarcinoma cell line, in which an auxin-inducible degron (AID) and a NeonGreen moiety were biallelically introduced in-frame at the endogenous NUP153 loci (Aksenova et al., 2020), we tracked lamin B2 ∼15 hours after auxin addition. During incubation with auxin, cells were either allowed to divide normally or kept in thymidine (added 8h before the time of auxin addition) to prevent progression into mitosis. Levels of Nup153 were confirmed to be similarly reduced under these conditions, as tracked using by immunoblot (Figure 6A). As DLD1 cells are not flat and tend to grow in clusters, identifying cells joined by a midbody structure was difficult to do with accuracy. We therefore identified individual nuclei with DNA stain and scored whether the corresponding lamin B2 was localized in foci rather than a peripheral nuclear rim. Auxin-mediated depletion of Nup153 robustly recapitulated the lamin B2 localization phenotype, with 53.6±5.4% of nuclei associated with lamin B2 in foci compared to 8.5±0.9% in vehicle treated cells (Figure 6B-C). Some focal lamin B2 was seen in control conditions; we noted that this was largely detected when the nuclei were in pairs, indicating they were newly formed sister cells (Figure 6B). The pattern of lamin B2 under these conditions was a qualitatively milder phenotype than following auxin-mediated Nup153 depletion and may reflect an intermediate step in nuclear assembly more evident in DLD1 cells. Lamin B2 levels remained similar in the absence or presence of Nup153 (Figure 6A). Importantly, when cell division was blocked in G1/S by thymidine treatment, focal lamin B2 was seen in very few cells – this phenotypic suppression was particularly striking in cells depleted of Nup153 (Figure 6C). Thus, the role for Nup153 is critical only when nuclear formation takes place (Figure 7).

**Figure 6.**
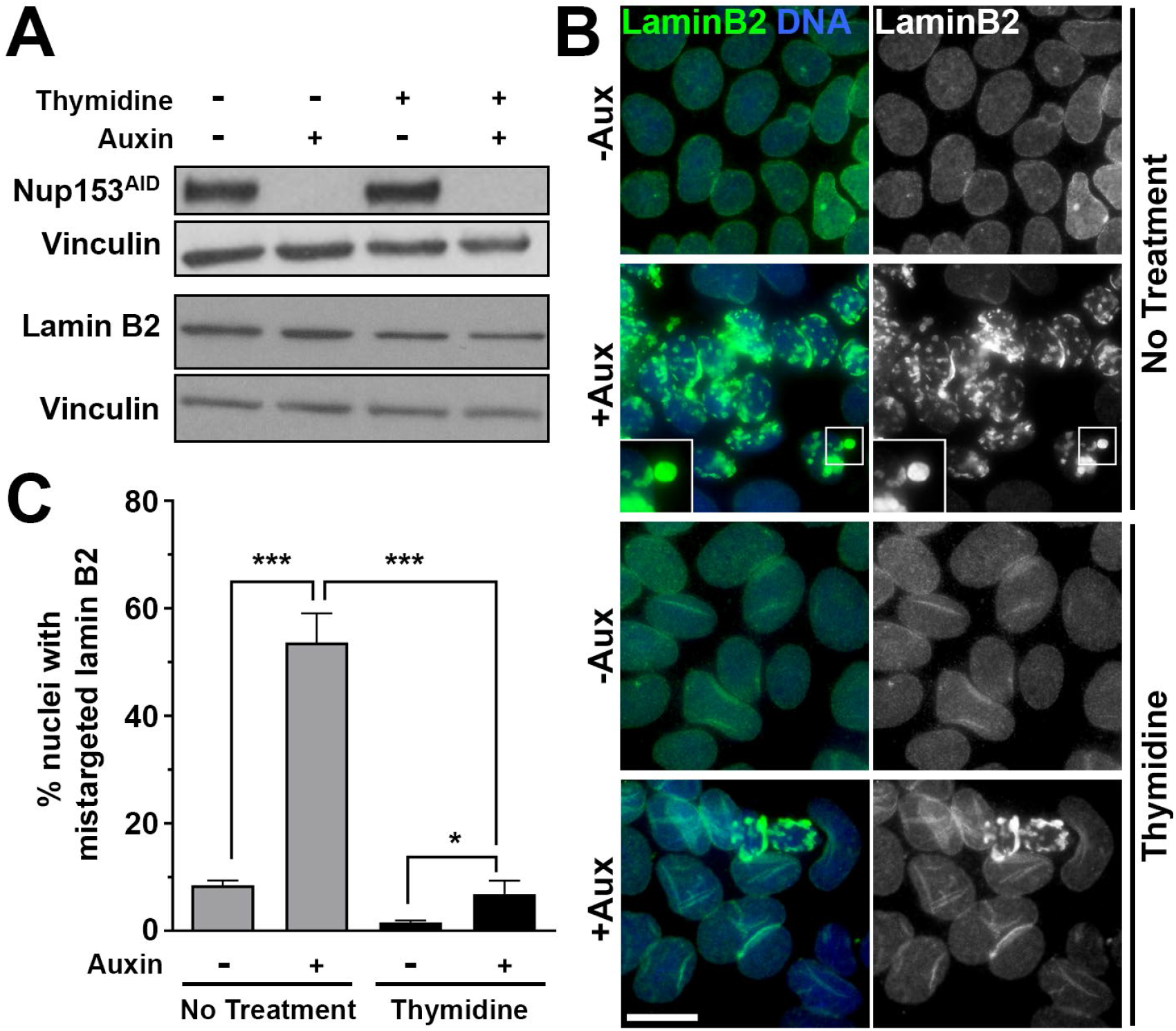
Nup153 plays a critical role specific to NE formation. (A) Immunoblot of DLD1-Nup153^AID^ cells growing asynchronously or in the presence of thymidine to prevent progression into mitosis. Cells were treated with auxin for 15 hours, as indicated. Following detection of Nup153 and lamin B2, blots were reprobed for vinculin as a loading control. (B) Widefield microscopy of cells treated as in A tracking the appearance of endogenous lamin B2 with DNA counterstained. Images were adjusted independently to best illustrate lamin B2 distribution pattern. Enlargements are 2X. Scale bar is 20 μM. (C) Quantification of nuclei with associated lamin B2 foci. Data collected from three biological replicates are graphed using the mean from each experiment to calculate the overall mean ± SD; no thymidine, no auxin 8.5±0.9% (N = 1078, 771, 784); no thymidine, plus auxin 53.6±5.4% (N = 664, 538, 465); thymidine, no auxin 1.6±0.4% (N = 486, 404, 389); thymidine, plus auxin 6.8±2.5% (N = 460, 353, 509). *P< 0.05, ***P<0.001.

**Figure 7.**
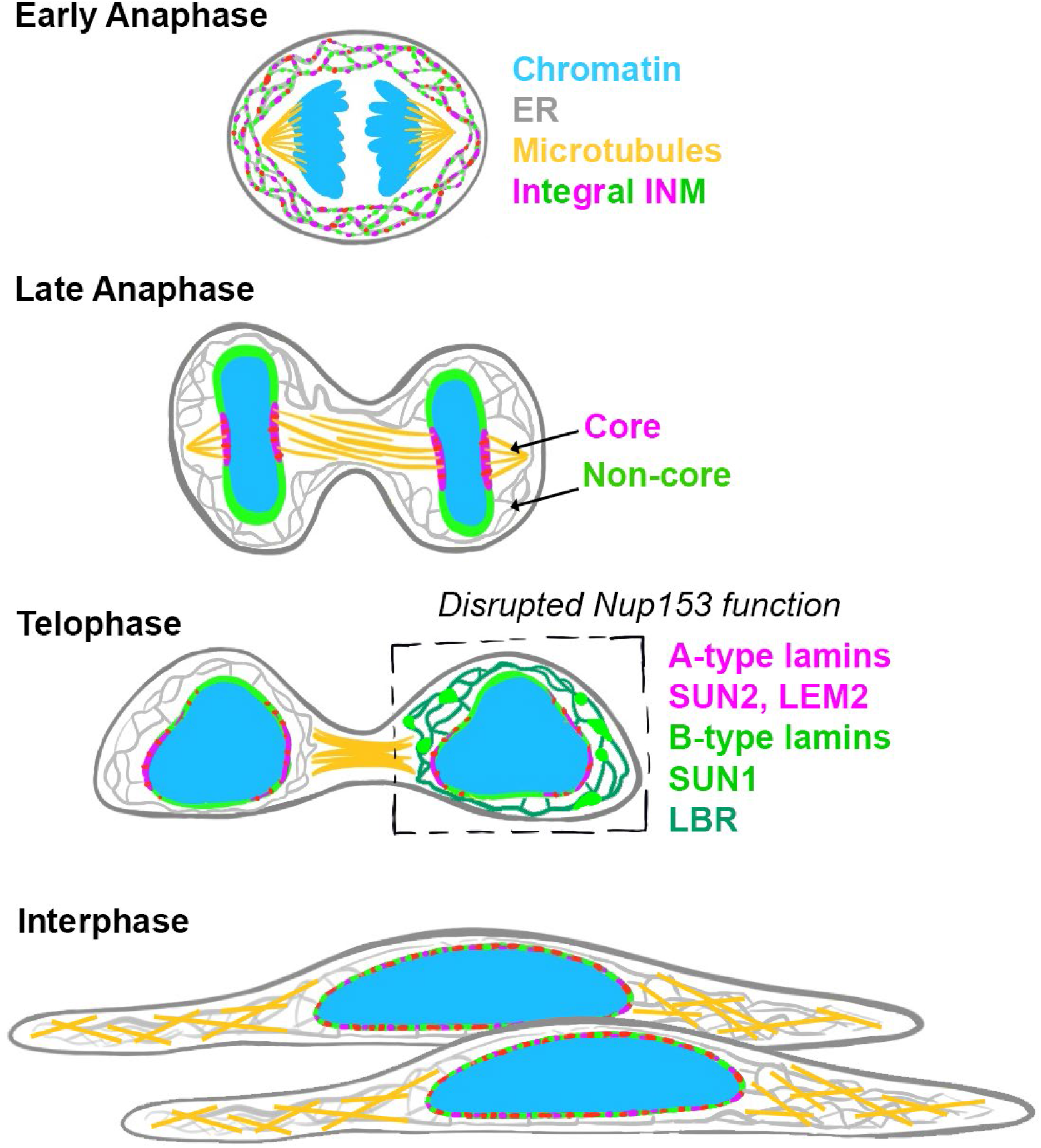
Summary model of observations. After the genome segregates in early anaphase, nuclear envelope proteins dispersed throughout the mitotic ER seed the formation of the nascent nuclear envelope at the chromatin surface. By late anaphase, INM proteins form distinct core and non-core domains at NE. As cells exit late anaphase and enter telophase, these transient domains redistribute as midzone microtubules condense into a midbody structure. During the progression from late anaphase to telophase, Nup153 is required for the continued targeting of primarily non-core INM proteins. At interphase, INM protein targeting occurs independently of Nup153.

## Discussion

### Nuclear envelope formation relies on Nup153

We previously probed the localization of lamin A following Nup153 depletion and concluded it was defective under this condition (Mackay et al., 2010). In following up on this observation, however, we found that the antibody we had employed, while marketed as an anti-lamin A antibody, was a pan-lamin antibody. Here, we used antibodies specific for each isoform and this revealed that the reliance on Nup153 is significantly more profound for B-type lamins. The specificity of this phenotype prompted us to consider membrane-associated proteins more broadly, and this inroad led to a new appreciation of a role for Nup153 in targeting a subset of integral membrane proteins to the INM during nuclear assembly. This phenotype was seen in three different cell lines (Hela, RPE1, and DLD1) and was elicited either by depletion of Nup153 (using two independent oligos or auxin-mediated degradation) or by expression of a dominant negative acting fragment of Nup153.

A similar dependence of B-type lamins on the nucleoporin ELYS has been reported (Jevtic et al., 2019), reinforcing the interconnected roles of these two Nups (Mimura et al., 2016). Yet, there are several distinctions between the contributions made by ELYS and Nup153 to nuclear assembly. First, ELYS has been shown to be crucial for early INM protein targeting events in anaphase that establish the core subdomain (Clever et al., 2012; Clever et al., 2013), whereas Nup153 appears dispensable for this aspect of nuclear assembly (Supplemental Figure 2). Second, ELYS plays a key role in post-mitotic NPC formation (Doucet et al., 2010; Franz et al., 2007; Rasala et al., 2006), while Nup153 is implicated in the ‘interphasic’ pathway of pore assembly (Vollmer et al., 2015). One possibility is that ELYS and Nup153 converge via a distinct and shared role to cooperatively contribute to lamin B localization. Alternatively, early defects in nuclear formation that arise from ELYS depletion may lead indirectly to aberrant B-type lamin targeting by preventing Nup153 from performing its function in this process at the time of NE expansion.

Persistence of LBR in the ER is a distinct phenotype that has also been observed when ELYS is depleted (Mimura et al., 2016). Notably, this phenotype arises very early in nuclear assembly and continues through interphase. The contribution of Nup153 to LBR targeting at interphase was reported previously (Mimura et al., 2016), and here we have found that this is also the case earlier, at a time coinciding with nuclear formation. In the case of ELYS, this phenotype has been attributed to its ability to interact directly with LBR and influence its phospho-regulation by the phosphatase PP1. Nup153 may participate in this pathway cooperatively or may independently reinforce this role.

Our finding that interference of Nup153, but not of Nup50 or Tpr, leads to a distinctive NE phenotype (Figure 2) is significant because it implicates Nup153 in having a role independent of its structural partners at the nuclear pore. Moreover, this finding uncouples roles that have been attributed to these other Nups and their contributions to parallel steps during NE formation. Such functions include a recently characterized role for Nup50 in regulating the Ran pathway via stimulation of its guanine nucleotide exchange factor RCC1 (Holzer et al., 2021). Future studies to elucidate the role of Nup153 and to parse the contribution of other nucleoporins during NE formation will be enabled by the information gained here on 1) the timing of dependence on Nup153 for NE protein targeting and 2) the identity of a specific subset of NE proteins dependent on Nup153 for targeting.

#### A multistage framework for nuclear assembly and maintenance

During nuclear formation, initial recruitment of NE residents to the main chromatin mass occurs relatively non-specifically, mediated by multiple protein-DNA/chromatin interactions attracting membrane to the chromatin surface (Anderson et al., 2009). Enrichment of core domain proteins rapidly ensues as they are attracted from the non-core periphery of the disk to concentrate at the central core region (Dechat et al., 2004; Haraguchi et al., 2008; Haraguchi et al., 2001). In the case of LEM2, we previously found that this recruitment relies on interactions with both the chromatin-associated protein BAF and spindle microtubules that remain attached to the disk at this time (von Appen et al., 2020). Following microtubule remodeling and ESCRT-mediated membrane closure, the sealed nuclear membrane must further expand to accommodate chromatin as it decondenses. How ongoing addition of proteins and lipids from the ER is accomplished is incompletely understood, but the results here indicate that Nup153 has a role in delivery of NE residents at this step. Specifically, our results identify a role for Nup153 as the NE expands following closure (Figure 7).

There are several noteworthy aspects about the requirement for Nup153 during nuclear envelope formation. First, we show that this role is important only for a subset of NE proteins, so it is not linked to a general expansion of the NE compartment. Second, those proteins affected by impairment of Nup153 function are primarily ones that are considered ‘non-core’ residents (LBR, SUN1, LAP2β, lamin B1, lamin B2). Thus, unexpectedly, even at a time when core and non-core domains no longer persist, these two categories of NE proteins differ in targeting requirements. It is likely too simplistic to view this as strictly a core/non-core dichotomy, but nonetheless these findings point to unexpected functional distinctions in NE protein targeting requirements at this step of nuclear assembly. Third, loss of Nup153 does not impact initial steps of nuclear formation, nor does it persist at interphase except in the case of LBR targeting. This suggests that building and maintaining the NE is a multi-phase process with distinct molecular steps mediating targeting of membrane and membrane-associated proteins.

This study provides a framework for understanding distinct facets of nuclear assembly and builds the groundwork for mechanistic dissection of the particular role(s) played by Nup153 during these steps. Nup153 has many attributes that may be key, including its ability to bind a wide range of partner proteins --e.g., transport factors, SUMO proteases, lamins, chromatin architectural proteins (Al-Haboubi et al., 2011; Chow et al., 2012; Hang and Dasso, 2002; Kadota et al., 2020; Nakielny et al., 1999; Vagnarelli et al., 2011; Zhang et al., 2002). The altered delivery of particular proteins to the NE may contribute further to the NE assembly phenotype. For instance, SUN1 has been implicated in chromatin decondensation and nuclear growth (Chi et al., 2007), thus its defective recruitment to the newly forming NE may alter the capacity for other proteins to recruit to this site.

As interference in Nup153 function also results in an active abscission checkpoint that controls the timing of the last step in cytokinesis (Mackay et al., 2010), it is tempting to speculate that the NE expansion step is under surveillance at this juncture. However, through the course of our experiments, we did not always see NE mislocalization phenotypes in cells halted with checkpoint activity at the midbody stage. Nonetheless, further studies are required to delineate whether NE protein targeting is required to satisfy the abscission checkpoint, as it is possible that a nuclear defect at this time does not always manifest as the full-blown phenotype tracked in this study.

#### Insights into the NE repertoire at parental and micronuclei

A theme that has been observed for micronuclei is their distinctive repertoire of NE proteins. This composition, which is often rich in core domain proteins and sparse in non-core proteins (Liu et al., 2018), leads to aberrant nuclear architecture and fragility and, in turn, a high potential for the collapse of NE integrity at micronuclei that is thought to contribute to genomic instability in tumors (Hatch et al., 2013; Liu et al., 2018; Zhang et al., 2015). Indeed, micronuclei lacking lamin B were specifically found to be prone to DNA damage (Kneissig et al., 2019). Our results here indicate that the low numbers of nuclear pore constituents, and low levels of Nup153 in particular (Hatch et al., 2013; Liu et al., 2018), at micronuclei may contribute to this pathological biogenesis in a way that was not previously recognized. Specifically, the role that Nup153 plays in ensuring non-core NE residents are recruited to nascent nuclei may be reduced at micronuclei and, at least in certain cases, there may not be an alternative route at interphase that allows this deficiency to recover, as we have seen happens in parent nuclei.

Notably, in cases where micronuclei form via a route that does not preclude non-core protein recruitment and NE expansion, the resulting NE are more stable and these micronuclei are not associated with tumorigenesis (Sepaniac et al., 2021). Such a situation is seen in KIF18A knockout mice, in which micronuclei arise following chromosome misalignment. It has been proposed that proximity of these micronuclei to ER membrane stores near the mitotic spindle may promote formation of stable NE. It will be of interest to determine whether recruitment of Nup153 is enhanced under these circumstances and plays a role in facilitating targeting of non-core residents to create a more robust NE. Ultimately, expanding our knowledge of requirements for INM protein recruitment throughout the cell cycle will clarify how cells maintain genomic stability and will lend insight into how this process goes awry at micronuclei.

## Materials and methods

### Cell culture and lines

HeLa cells were a gift from Maureen Powers and RPE-1 cells were a gift from Bruce Edgar (RRID:CVCL 4388) and the identity of both cell types were confirmed by STR profiling. HeLa and RPE-1 cells were grown in DMEM supplemented with 10% fetal bovine serum at 37°C with 5% CO_2_. RPE-1 cells are immortalized by the presence of hTERT, expression of which was maintained by continued hygromycin treatment. HeLa cells stably expressing GFP-lamin B2 and LEM2-mCherry together were established previously (von Appen et al., 2020). HeLa cells stably expressing H2B-mCherry were established previously (Mackay et al., 2010) and used to generate cells co-expressing the GFP-lamin B2 construct from Bob Goldman. Cells expressing BFP-Sec61β were generated from a construct from Gia Voeltz obtained through Addgene (plasmid #49154; http://n2t.net/addgene:49154; RRID:Addgene_49154) (Zurek et al., 2011). Stable cell lines were established by transfecting plasmids using Lipofectamine 2000 or Lipofectamine 3000 (Invitrogen), as per the manufacturer’s instructions, and selecting for puromycin or G418 resistance. Single colonies were expanded and screened for expression. Cell lines stably expressing Tet-inducible GFP or GFP-tagged Nup153-C were established and protein expression was induced as previously described (Mackay et al., 2010). DLD1 cells were established previously and maintained as described (Aksenova et al., 2020).

### Auxin-mediated Nup153 degradation

DLD1-Nup153^AID^ cells were plated at 140K per well on glass coverslips and cultured for 8 hours in the absence or presence of 5 mM thymidine (Calbiochem). 1 mM Indole-3-acetic acid (IAA, referred to here as auxin; Sigma-Aldrich) was then added to half of the cells from each condition and cells were cultured an additional 15 hours, continuing with and without thymidine. Cells were fixed for immunofluorescence. Parallel cultures in 6-well dishes seeded with 700K cells were treated identically and whole cell lysates for immunoblot.

### RNA interference

Hela or RPE-1 cells were transfected with Lipofectamine RNAiMAX (Invitrogen) and final concentration of 10 nM siRNA oligonucleotides: siCon, GCAAAUCUCCGAUCGUAGA (Mackay et al., 2009), si153, GGACUUGUUAGAUCUAGUU, (Mackay et al., 2009), si153-1 (Harborth et al., 2001), si50 (Ogawa et al., 2010), siTpr (Mackay et al., 2010), where indicated. Cells were assessed 48 hours after transfection for asynchronous experiments.

### HeLa cell synchronization

At the time of plating or eight hours after plating cells with siRNA transfection mixture, 2mM thymidine was added to the cells without changing medium containing the transfection mixture. After 24 hours, cells were rinsed thoroughly three times with PBS before exchanging with complete culture medium without transfection mixture. To track the kinetics of release from G1/S arrest, cells were assessed 10 and 16 hours after thymidine wash-out. All other synchronization experiments assessing Nup153-depletion phenotypes were fixed 16 hours following thymidine wash-out.

### Microscopy

For live cell imaging, cells were plated with siRNA transfection mixture on untreated 25mm glass-bottomed MatTek dishes. After 48 hours, medium was exchanged for phenol-free imaging medium supplemented with 10% FBS. Mitotic cells were imaged using spinning disk confocal microscopy at 37°C with 5% CO_2_ on a Nikon Eclipse T*i* microscope with a 60x objective. Images were acquired using Metamorph software. Cells stably expressing GFP-lamin B2 and H2B-mCherry were imaged every 3 minutes. Cells stably expressing GFP-lamin B2 and LEM2-mCherry were imaged 1 hour following the addition of NucBlue (Invitrogen), per manufacturer’s instructions, and imaged every 15s. Fixed imaging was performed by spinning disk confocal microscopy (Nikon Eclipse T*i*; Metamorph) or by widefield microscopy (Zeiss Axioskop 2 with 63x objective; Zeiss Axiovision and Zeiss Zen Blue).

### Data analysis

Plot line profile analysis was performed on raw images acquired by spinning disk confocal microscopy using the ‘Plot Profile’ function in ImageJ.

For comparison of SUN1 and SUN2 enrichment at ectopic foci relative to the nuclear envelope following Nup153-depletion, nuclei were defined as ROIs using DAPI while ectopic SUN1 was used to define ROIs characteristic of the Nup153-depletion phenotype using Zeiss Zen Blue semi-automated Image Analysis module. The ratio of mean fluorescence (arbitrary units) of ectopic to nuclear SUN1 and SUN2 were calculated and plotted from three independent experiments.

To obtain the cytoplasmic to nuclear LBR ratio in cells treated with either siRNA targeting Nup153 or control siRNA, the fluorescence intensity of LBR in the nucleus (identified with DAPI staining) and in the cytoplasm of midbody stage (identified using tubulin staining) cells was measured using ImageJ. The fluorescence intensity at the nucleus was measured using a circle 5 microns in diameter placed at the center of each nucleus. The intensity at the cytoplasm was measured using two circles 5 microns in diameter placed on opposite sides of the midbody. The mean intensities were measured and used to obtain the cytoplasmic to nuclear LBR ratios for each cell. Only Nup153-depleted cells that had ectopic LaminB2 foci were quantified.

All plots generated using GraphPad Prism.

### Immunofluorescence

Cells were plated on untreated coverslips for experimental manipulation and fixed in either −20°C methanol for 5-10 minutes or 2% paraformaldehyde for 15 minutes. Primary and secondary antibody incubations were performed in blocking buffer for 1 hour at room temperature. Primary antibodies are as follows: rabbit α-Aurora B pT232 (Rockland 600-401-677); mouse α-lamin A/C (Cell Signaling 4C11); rabbit α-lamin B1 (Abcam ab16408); mouse α-lamin B2 (Abcam ab8983); hamster α-Lap2α (gift from Joe Glavy); mouse α-Lap2β (gift from Brian Burke); rabbit α-LBR (Abcam ab32535); mouse α-Nup153 (SA1; gift from Brian Burke); rabbit α-POM121 (GeneTex GTX102128); rabbit α-SUN1 (Abcam ab124770); mouse α-SUN2 (gift from Brian Burke); rabbit α-Tubulin (Abcam ab18251); rat α-Tubulin (Abcam ab6160); chicken α-Tubulin (Synaptic systems; 302 206)] followed by AlexaFluor conjugated secondary antibodies (ThermoFisher). Coverslips were mounted on slides using DAPI Prolong Gold, per manufacturer’s instructions (Invitrogen).

### Immunoblotting

Cells were plated in 6-well dishes and experimentally manipulated in parallel with immunofluorescence experiments where applicable. Cells were harvested by scraping and pelleting by centrifugation. DLD1 lysates were prepared in RIPA buffer (150 mM NaCl, 0.5% NP-40, 0.1% SDS, 50 mM Tris-HCL pH 8,100 mM NaF, 0.2 mM Sodium orthovanadate) and HeLa cell lysates were prepared in NP40 lysis buffer (250 mM NaCl, 1% NP-40, 50 mM Tris pH 7.4, 5 mM EDTA, 50 mM NaF, 1 mM Sodium orthovanadate). Lysates were cleared by centrifugation and quantified by Pierce Coomassie Bradford Assay Reagent (ThermoFisher). Samples were prepared at equal concentrations in SDS loading buffer, boiled, and resolved by SDS-PAGE. Following wet transfer, PVDF membranes were blocked in 3% milk in TBS-T (0.1% Tween-20 in Tris-buffered saline). Primary and secondary antibody incubations were also performed in 3% milk in TBS-T at room temperature for 1 hr each. For enhanced chemiluminescence detection, secondary antibodies conjugated with HRP (ThermoFisher) were detected using Western Lightning PLUS ECL (PerkinElmer, Waltham, MA) on Hyblot CL film (Thomas Scientific, Swedesboro, NJ). Alternatively, secondary antibodies conjugated with near-infrared fluorescent dyes (Abcam, Cambridge, UK) were detected using a Li-Cor Odyssey Infrared scanner.

Lysates from HeLa cells transiently expressing GFP-lamin fusions (gift from Bob Goldman) following transfection with Lipofectamine LTX (ThermoFisher) were used to test lamin antibody specificity by probing immunoblots with mouse α-lamin A/C (Cell Signaling 4C11), mouse α-lamin B1 (Abcam ab16408), mouse α-lamin B2 (Abcam ab8983), and rabbit α-GFP (Abcam ab290). Nucleoporin depletion was confirmed using the indicated antibodies: mouse α-Nup153 (SA1; gift from Brian Burke), rabbit α-Nup50 (custom antibody (Mackay et al., 2010)), rabbit α-Nup155 (Genetex GTX120945), and rabbit α-Tpr; Bethyl Laboratories IHC-00099-1). DLD1 lysates were probed for Nup153 (SA1), mouse α-lamin B2 (Abcam ab8983), α-vinculin (Sigma V9131).

## Abbreviations

AID: auxin inducible degradation
DAPI: 4′,6-diamidino-2-phenylindole
ER: endoplasmic reticulum
ESCRT: endosomal sorting complexes required for transport
INM: inner nuclear membrane
LBR: lamin B receptor
NE: nuclear envelope
NPC: nuclear pore complex
SUN1/2: Sad1 and UNC84 Domain Containing

## Acknowledgements

We would like to thank Suzanne Elgort for her contributions to the initiation of this project and Dr. Michelle Mendoza for providing advice as well as access to a spinning disk confocal microscope. We thank Drs. Bob Goldman, Joe Glavy, and Brian Burke for sharing reagents, as indicated in Materials and Methods. This work was supported by R01GM131052 (KSU) and an Allen Distinguished Investigator Award, a Paul G. Allen Frontiers Group advised grant of the Paul G. Allen Family Foundation (MN and KSU).

## Notes

### Competing Interest Statement

The authors have declared no competing interest.

