## Supplemental Figures for "A Role for Nup153 in Nuclear Assembly Reveals Differential Requirements for Targeting of Nuclear Envelope Constituents"

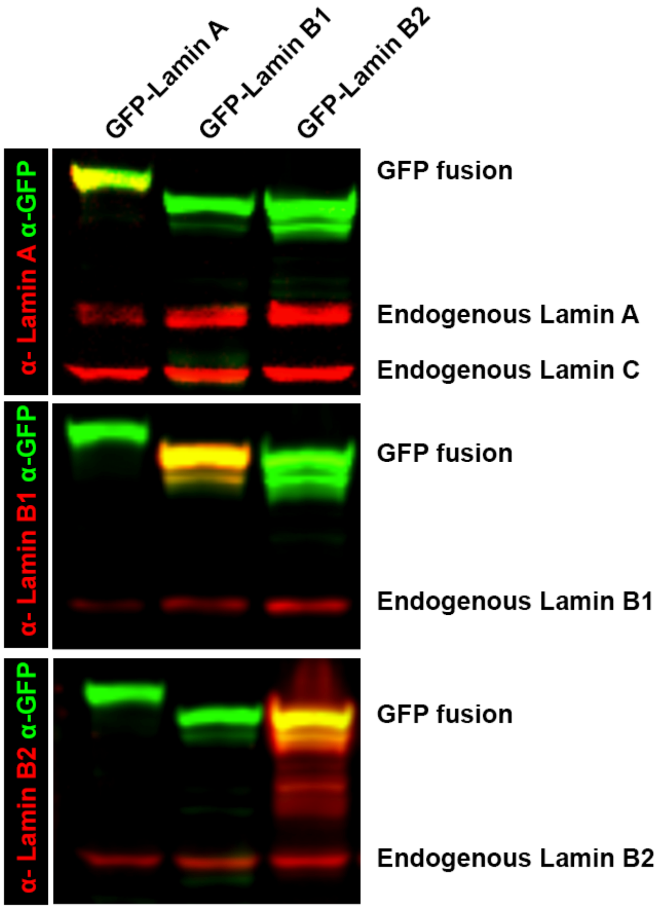


**Supplemental Figure 1. Lamin antibodies are specific to individual lamin proteins.** Lysates from HeLa cells transiently transfected with expression constructs for either GFP-Lamin A, GFP-Lamin B2, GFP-Lamin B2, or GFP alone were probed using α-Lamin A (Cell Signaling 4C11), α-Lamin B1 (Abcam ab16408), α-Lamin B2 (Abcam ab8983), and α-GFP (Abcam ab290).

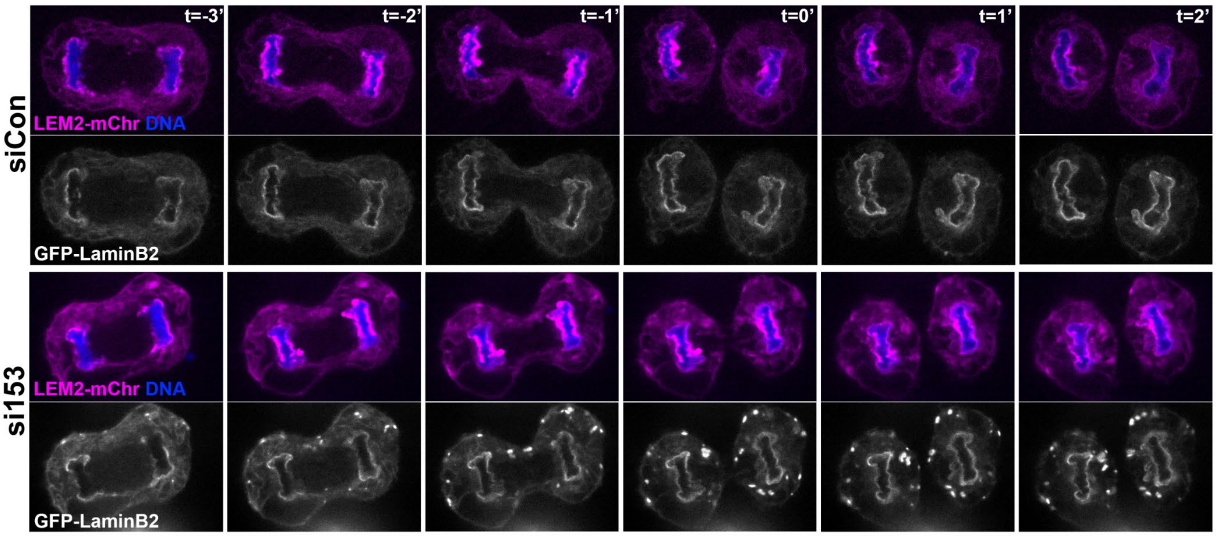


**Supplemental Figure 2. Nuclear enclosure and dynamic formation of the core domain of the NE occur in Nup153 depleted cells.** Live cell imaging by spinning disk confocal microscopy showing LEM2-mCherry clustering at the core domain of nascent nuclei in both siCon and si153 cells. In parallel, GFP-Lamin B2 targeting is shown. T=0' indicates the time of complete cleavage furrow ingression.

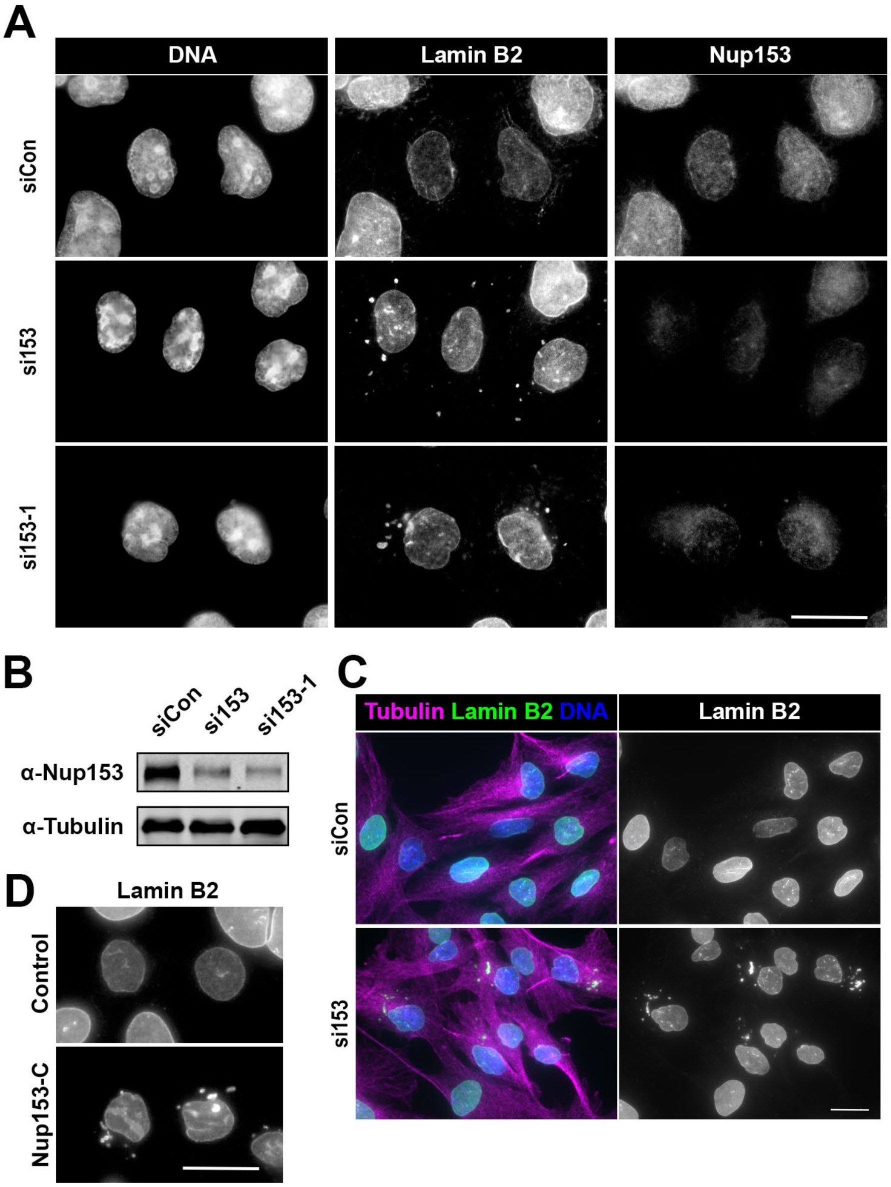


**Supplemental Figure 3. The Nup153 depletion phenotype is recapitulated in different experimental scenarios.** (A) Representative images of cells stained for Nup153 and lamin B2 by widefield microscopy after 48-hour treatment with the indicated siRNAs. (B) Immunoblot of Nup153 and tubulin, as a loading control, after 48-hour treatment with the indicated siRNAs. (C) Representative images of RPE-1 cells treated with siCon or si153 for 48 hours and stained for lamin B2 and tubulin. (D) Control cells or those expressing the Nup153-C fragment stained for lamin B2 and imaged by widefield microscopy. All scale bars 20μm.

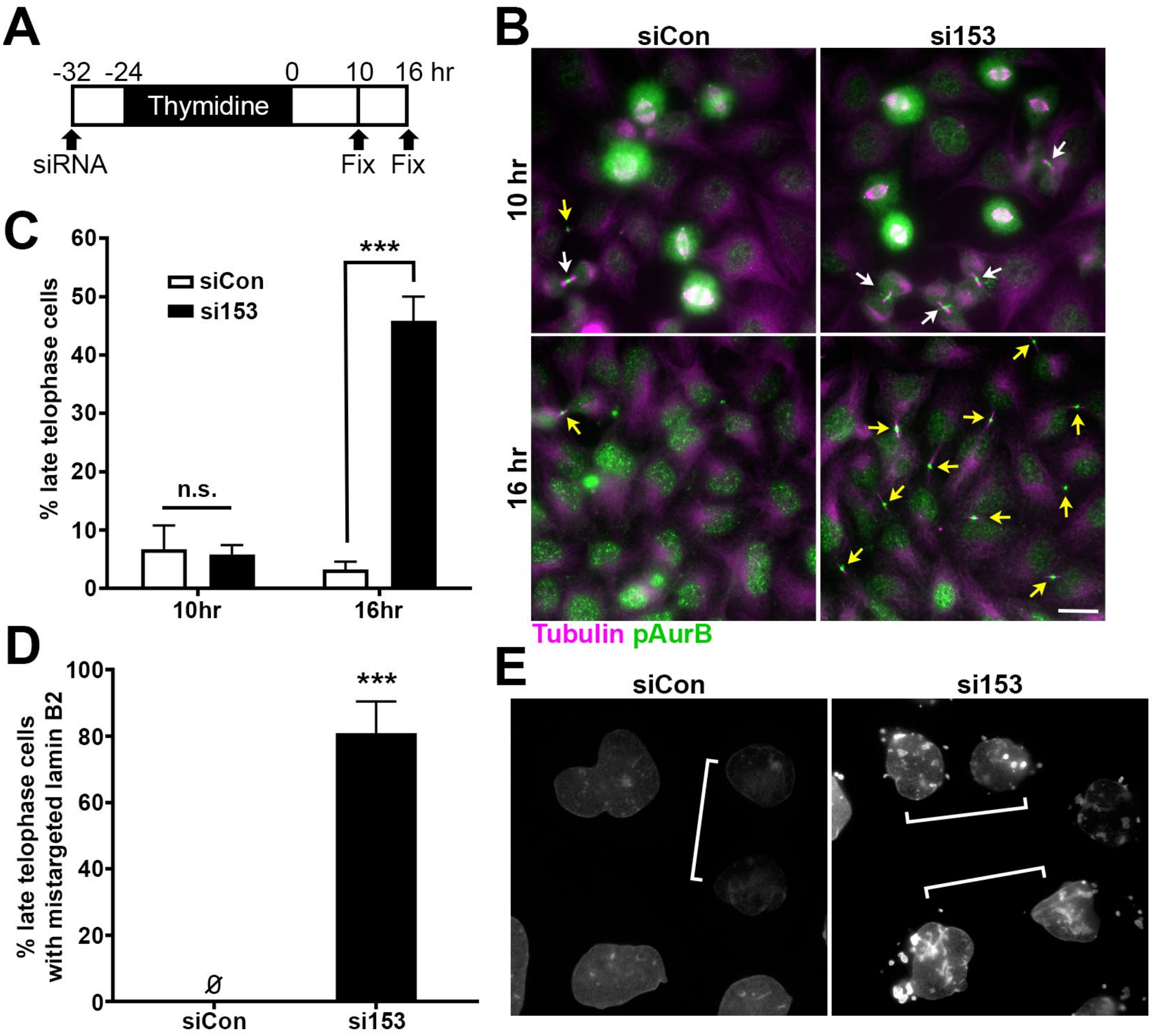


**Supplemental Figure 4. Synchronization enriches for late telophase-stage cells.** (A) Timeline of synchronization experiment. (B) Widefield images of cells treated with siCon or si153 after 10 and 16 hours thymidine release, stained for phospho-Aurora B and tubulin. Scale bar 20µm. (C) Quantification of late telophase cells after treatment with siCon or si153 after 10 and 16 hours after thymidine release. Mean ±SD; siCon 10hr: 7±4%, n=3; si153 10hr: 6±2%, n=3; siCon 16hr: 3±1%, n=4; si153 16hr: 45±4%, n=4; n.s. = not significant, ***P < 0.001. (D) Quantification of the percent late telophase-stage cells with mistargeted lamin B2 after treatment with siCon or si153. Mean ±SD; siCon: 0±0%, n=4; si153: 81±10%, n=4; ***P < 0.001. Ø indicates raw values were equal to 0. (E) Representative images of lamin B2 in late telophase cells (as indicated by tubulin co-stain, not shown), denoted by white brackets, in control and Nup153-depleted cells.

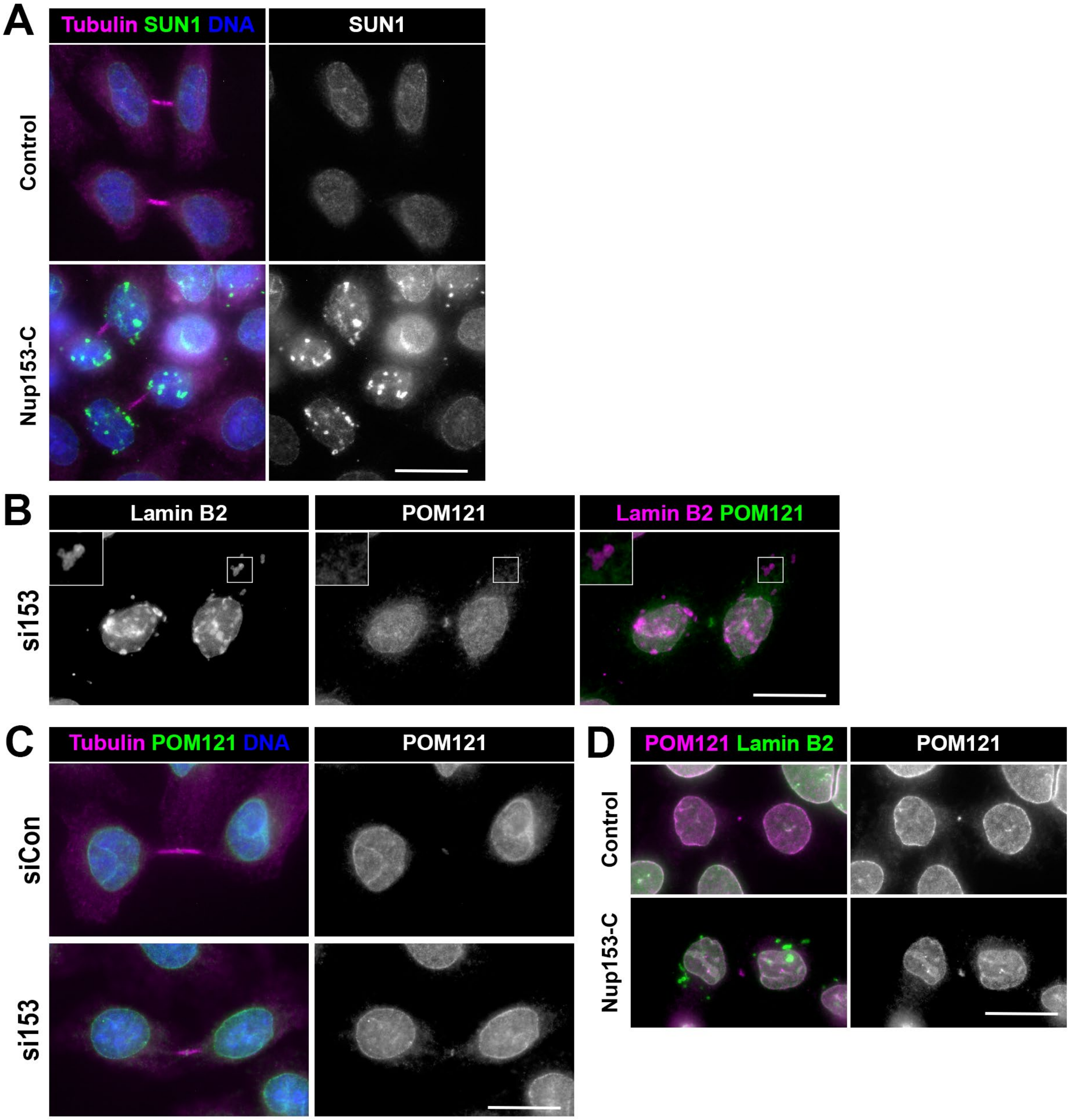


**Supplemental Figure 5. Further analysis of integral membrane proteins of the nuclear envelope.** (A) Representative images of control and Nup153-C-expressing cells stained for tubulin and SUN1, imaged by widefield microscopy. (B) Representative image of lamin B2 and POM121 in Nup153-depleted and thymidine-synchronized cells. Enlargements are 2x. (C) Representative images of POM121 in late telophase-stage cells after siCon and si153 treatment and thymidine synchronization. (D) Representative images of control and Nup153-C-expressing cells stained for lamin B2 and POM121, imaged by widefield microscopy. All scale bars 20μm.

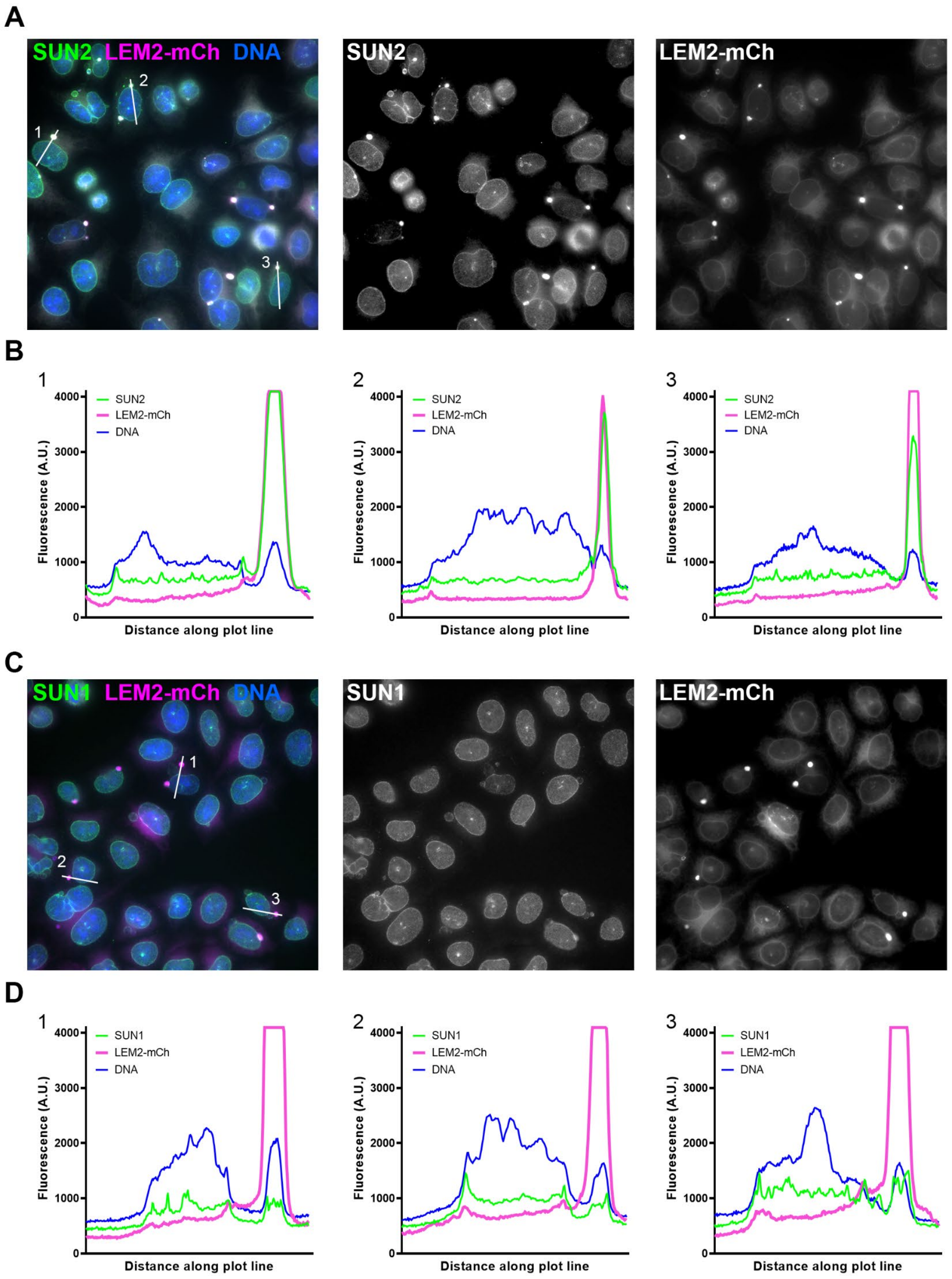


**Supplemental Figure 6. SUN2, but not SUN1, enriches at core-enriched micronuclei.** (A) Wide-field images of endogenous SUN2 detection alongside LEM2-mCherry. (B) Arbitrary fluorescence units corresponding to plot lines shown in A are graphed. (C) Wide-field images of endogenous SUN1 detected alongside LEM2-mCherry. (D) Arbitrary fluorescence units corresponding to plot lines shown in C are graphed.
